# Delayed primacy recall performance predicts *post mortem* Alzheimer’s disease pathology from unimpaired *ante mortem* cognitive baseline

**DOI:** 10.1101/2023.06.26.546225

**Authors:** Davide Bruno, Kristina M. Gicas, Ainara Jauregi Zinkunegi, Kimberly D. Mueller, Melissa Lamar

## Abstract

**INTRODUCTION:** We propose a novel method to assess delayed primacy in the CERAD memory test. We then examine whether this measure predicts *post mortem* Alzheimer’s disease (AD) neuropathology in individuals who were clinically unimpaired at baseline.

**METHODS:** A total of 1096 individuals were selected from the Rush Alzheimer’s Disease Center database registry. All participants were clinically unimpaired at baseline, and had subsequently undergone brain autopsy. Average age at baseline was 78.8 (6.92). A Bayesian regression analysis was carried out with global pathology as outcome; demographic, clinical and *APOE* data as covariates; and cognitive predictors, including delayed primacy.

**RESULTS:** Global AD pathology was best predicted by delayed primacy. Secondary analyses showed that delayed primacy was mostly associated with neuritic plaques, whereas total delayed recall was associated with neurofibrillary tangles.

**DISCUSSION:** We conclude that CERAD-derived delayed primacy is a useful metric for early detection and diagnosis of AD in unimpaired individuals.

## 1. Background

Promoting early detection and diagnosis of Alzheimer’s disease (AD) is one of the critical components of the global response to the growing dementia crisis [1]. A timely diagnosis of AD can promote patients’ safety, help avoid preventable hospitalizations, aid identification of caregivers, and support financial planning [2,3,4]. In addition, early detection of AD can facilitate the selection of individuals for clinical trials [5].

Research into the use of biomarkers for early detection and diagnosis of AD has seen substantial progress in recent years, including the introduction of promising blood-based biomarkers (see e.g., [6]). Despite this, many studies still rely in great part on assessments based upon positron-emission tomography (PET) imaging and/or lumbar puncture. These tests can be intimidating, and require access to highly specialized clinical settings [7]. As the burden of dementia worldwide is affecting especially low and middle income countries [8], early detection frameworks should focus on accurate, but also affordable and accessible technologies.

Testing neuropsychological function is non-invasive, requires relatively minimal training, and is inexpensive. Of relevance to AD is especially the assessment of episodic memory ability [9,10,11,12]. However, neuropsychological test metrics were designed primarily for diagnostic purposes, i.e., the identification of well-defined changes in performance, and not for the detection of subtle shifts in neuropsychological function due to emergent underlying pathology [13]. An alternative to the examination of standard clinical metrics is process analysis of neuropsychological test performance (or the Boston process approach; [14,15]).

Analysis of process scores is based upon the principle that different cognitive processes underlie overall test performance, and that unearthing these processes may be more informative than simply evaluating typical composite scores.

Examples of effective process scores applied to verbal memory testing derive from the analysis of serial position performance. The serial position curve is a common pattern in tests of human memory, where performance tends to be better for stimuli learned at the beginning (primacy) and/or at the end (recency) of a list, as compared to those in the middle (hence the curve shape; e.g., Murdock, 1962). This recall pattern has been reproduced countless times. Primacy effects have been ascribed to extra opportunities for rehearsal [17], edge effects [18], and increased attention [19], among other things, whereas recency effects have been associated primarily to working memory processing [20]. Importantly, participants with AD diagnoses present with specific patterns of serial position performance, and these patterns can be examined for the purposes of early detection and diagnosis. In particular, a reduction of the primacy effect has been reported frequently in immediate recall tests in conjunction with AD risk and pathology (e.g., [21,22]). Gicas et al. [23], for example, have shown that higher immediate primacy performance associated with less risk of AD neuropathology, including hippocampal sclerosis; and increased pathology predicted greater longitudinal decline in primacy peformance [24].

Gicas et al. [23,24] measured serial position performance using the Consortium to Establish a Registry for Alzheimer’s Disease (CERAD) memory test. In this memory test, participants are presented with a list of ten words three times, in different presentation orders. Immediately after each presentation, participants are asked to free recall the words in any order (immediate recall). Subsequently, they are also asked to free recall the words after a delay (delayed recall). So far, analysis of serial position effects in the CERAD memory test has focused exclusively on immediate word-list recall from the first list, as the presentation order of the words changes in each learning trial. However, past analyses of word-list learning performance with tests other than CERAD have shown delayed primacy to be a better longitudinal predictor of global cognitive decline [25], and mild cognitive impairment (MCI; [26]) compared to immediate primacy. A reason for this may be that delayed primacy is sensitive to synaptic consolidation, as suggested also by its association with hippocampal grey matter volume [27] and increased functional connectivity between left and right hippocampus [28]. Therefore, the aim of the present study was to determine whether delayed primacy in CERAD was more effective at predicting AD neuropathology than immediate primacy.

In order to extract delayed primacy information from CERAD, we propose that each time the word list (A, B and C) is presented, one item will be shown *first*. The three items shown first (i.e., A1, B1, and C1) should all benefit from primacy exposure (e.g., extra opportunity for rehearsal, edge effects, etc…). Hence, when examining delayed recall performance, memory for items A1, B1 and C1 will give an account of delayed primacy. Using this method, we carried out secondary data analyses on Rush Alzheimer’s Disease Center cohort data. We hypothesized that a measure of delayed primacy, extracted from the CERAD memory test, would be effective in predicting *post mortem* AD-related pathology from a cognitively unimpaired baseline.

## 2. Methods

### 2.1 Participants

A total of 5158 unique participants’ data were available from the Rush Alzheimer’s Disease Center (RADC) database registry. From this total, we selected individuals who a) were diagnosed as having no cognitive impairment at baseline (see below for diagnostic criteria); b) had received a *post mortem* examination; c) were diagnosed as being cognitively unimpaired, or having MCI or Alzheimer’s disease at death (i.e., mixed cases were excluded; see below for diagnostic criteria); d) had the necessary CERAD data at baseline; and e) had *APOE* genotype data. Once these criteria were applied, the overall sample size was reduced to 1096. These participants came from four different RADC cohort studies: The Religious Order Study (ROS; n=486); the Memory and Aging Project (MAP; n=577); the Minority Aging Research Study (MARS; n=32); and the Latino Core Study (LATC; n=1). ROS [29] started in 1994 and comprises 65 year and older Catholic nuns, priests and brothers from across the USA. All were without known dementia at enrolment, and agreed to yearly evaluation and brain donation after death. MAP started in 1997 and comprises 65 year and older adults from retirement communities and subsidized senior housing in the Chicago area, and North-Eastern Illinois. All were without known dementia at enrolment, and agreed to yearly evaluation and brain donation after death. MARS [30] started in 2004 and is made up of 65 year and older adults identifying as African American and living in the Chicago area and suburbs. All were without known dementia at enrolment, and agreed to yearly evaluations, in addition to optional brain donation after death. LATC [31] started in 2015 and also includes 65 year and older adults from the Chicago area, who identify as Latino/Hispanic. All were without known dementia at enrolment, and agreed to yearly evaluations, in addition to optional brain donation after death. Overall, average age at baseline was 78.8 (6.93), average time between baseline and death was 10.56 (5.90) years, and average years of education were 16.29 (3.72). In total, 776 (71%) were female, and the *APOE* genotype was distributed as follows: four had ε2ε2; 151 had ε2ε3; 26 had ε2ε4; 711 had ε3ε3; 191 had ε3ε4; and 13 had ε4ε4. All participants signed a Repository Consent to allow their data to be shared. Ethics approvals for the studies included in these analyses were obtained from an Institutional Review Board of Rush University Medical Center. This research was completed in accordance with the Helsinki Declaration.

### 2.2 Baseline diagnosis

[32,33] was based on a three-stage process including computer scoring of a cognitive battery of 19 cognitive tests, clinical judgment by a neuropsychologist blinded to participants demographics, and a final diagnostic classification by a clinician (neurologist, geriatrician, or geriatric nurse practitioner). Clinical diagnosis of Alzheimer’s dementia is based on criteria of the joint working group of the National Institute of Neurological and Communicative Disorders and Stroke and the Alzheimer’s Disease and Related Disorders Association (NINCDS/ADRDA). The diagnosis of Alzheimer’s disease requires evidence of a meaningful decline in cognitive function relative to a previous level of performance with impairment in memory and at least one other area of cognition. Diagnosis of MCI is rendered for persons who are judged to have cognitive impairment by the neuropsychologist but are judged not to meet criteria for dementia by the clinician. Finally, persons without dementia or MCI are categorized as having no cognitive impairment.

### 2.3 Diagnosis at death

was based on assessment of all available clinical data, which was reviewed by a neurologist with expertise in dementia, leading to a summary diagnostic opinion rendered regarding the most likely clinical diagnosis at the time of death. Summary diagnoses were blind to all *post mortem* data. Case conferences including one or more neurologists and a neuropsychologist were used for consensus on selected cases.

### 2.4 Cognitive assessment

Participants completed a 19-test neuropsychological evaluation in a standardized format to assess functioning in the domains of episodic memory, semantic memory, working memory, visuospatial ability, and perceptual speed [34]. The raw scores from these tests were then converted to z-scores and averaged to create a global cognitive function index. The CERAD Word List Memory test [35] was part of this battery, and was the basis for the serial position scores. This test is composed of a list of 10 semantically unrelated words that are repeated across three trials (A, B and C) with varying word order. Participants are asked to recall as many words as possible immediately after presentation of each word list. Performance over the three trials is combined to give a total recall score. Later, after a delay of several minutes, participants are asked to recall as many words from the initial list as possible in any order, providing a total delayed recall score. Following Gicas et al. [23,24] we defined *immediate primacy* as the ability to recall the first three words presented in the first three trials (i.e., A1-A3, B1-B3, and C1-C3), immediately after presentation of that list, and then divided that sum by the total number of possible items (i.e., nine). *Delayed primacy* was defined as the ability to recall the first word presented in the first trial, the first words presented in the second trial, and the first word presented in the third trial (i.e., A1, B1, and C1), in the delayed recall task.

### 2.5 Post mortem evaluation

All participants in these analyses underwent brain autopsy at death [33]. The standard protocol involved removal of the brainstem and cerebellum, followed by cutting one hemisphere into 1cm coronal slabs that were immediately frozen. The other hemisphere was fixed in 4% paraformaldehyde for 3 to 21 days and subsequently cut into 1 cm coronal slabs. Regional blocks of tissue were embedded in paraffin, sliced into 6 μm sections, and mounted to glass slides for microscopic evaluation by a neuropathologist blinded to all clinical information.

A global AD pathology burden score was then extracted as a quantitative summary of AD pathology. This score derived from the average of three AD pathologies: neuritic plaques, diffuse plaques, and neurofibrillary tangles, as determined by microscopic examination of silver-stained slides from 5 regions: midfrontal cortex, midtemporal cortex, inferior parietal cortex, entorhinal cortex, and hippocampus.

### 2.6 APOE genotyping

DNA was extracted from PBMCs or brain. Participants were genotyped for APOE alleles by Polymorphic DNA Technologies (http://www.polymorphicdna.com/).

### 2.7 Data analysis

Bayesian regression analyses were carried out. In these analyses, all combinations of predictors were tested for data fit, including models with only one predictor, against the null model. The primary analysis had global AD pathology as outcome; *APOE* genotype, baseline age, sex, years of education, final diagnosis, and time from baseline to death as control variables (null model); and cognitive scores as predictors: total recall, total delayed recall, immediate primacy, and delayed primacy were derived from CERAD, while the global cognitive function index was obtained from the test battery (see Cognitive Assessment). Credible intervals (CIs) were set to 95%. The prior was set to JZS, and the model prior was set to Uniform. One thousand Markov chain Monte-Carlo simulations were conducted to determine parameters. Secondary analyses were carried out with neuritic plaques, diffuse plaques, and neurofibrillary tangles as outcomes, while otherwise maintaining the same parameters as in the primary analysis. Analyses were carried out in JASP 0.17.1.

## 3. Results

Table 1 reports means and standard deviations for the neuropathological and memory scores. Global AD pathology was best predicted by delayed primacy, BF_10_ = 5.577, with moderate evidence. This model was roughly twice as effective as the second best model (BF_10_ = 2.681) including total delayed recall only. The mean coefficient for delayed primacy was -0.028 (0.028; 95% credible intervals from -0.077 to 0), indicating that more global AD pathology was associated to poorer delayed primacy at baseline (Figure 1). These numbers suggest that, comparing cross-sectionally, one *less* unit of primacy corresponded to ∼ 4.2% *more* global AD pathology (based on the mean overall level of global AD pathology; Table 1): hence, comparing a person with 0 delayed primacy to a person with full delayed primacy (i.e., 3 units), the former will have upwards of 12% more global AD pathology. BF_inclusion_ for delayed primacy was 1.467, suggesting that adding delayed primacy to the model improved its fit to the data by 47%.

**Table 1.**
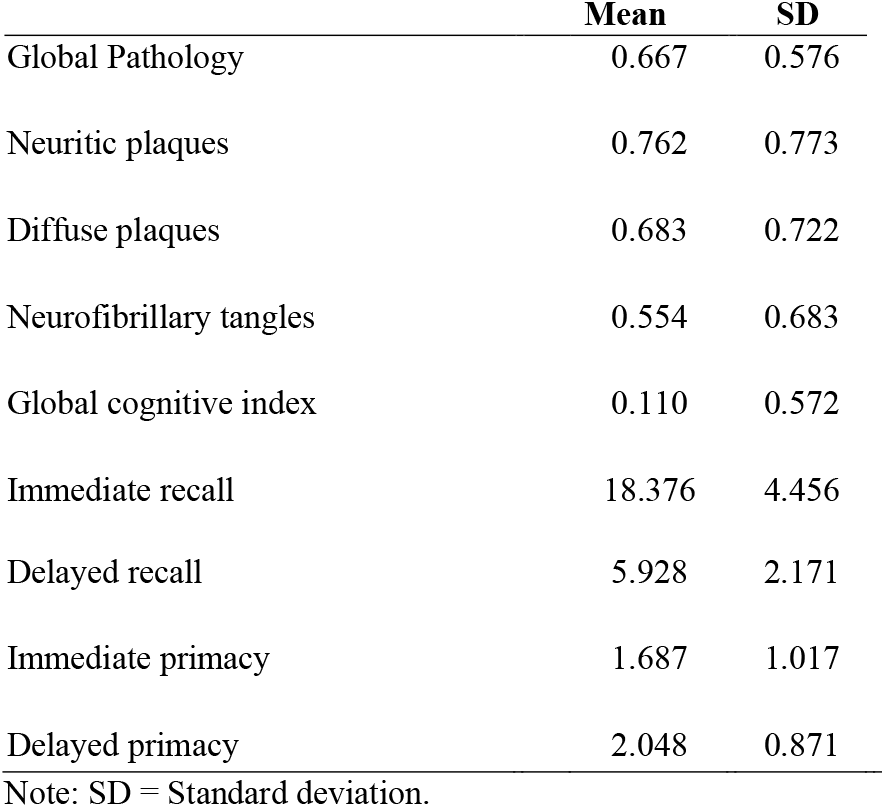
Neuropathological and memory data.

|  | **Mean** | **SD** |
| --- | --- | --- |
| Global Pathology | 0.667 | 0.576 |
| Neuritic plaques | 0.762 | 0.773 |
| Diffuse plaques | 0.683 | 0.722 |
| Neurofibrillary tangles | 0.554 | 0.683 |
| Global cognitive index | 0.110 | 0.572 |
| Immediate recall | 18.376 | 4.456 |
| Delayed recall | 5.928 | 2.171 |
| Immediate primacy | 1.687 | 1.017 |
| Delayed primacy | 2.048 | 0.871 |

Note: SD = Standard deviation.

**Figure 1.**
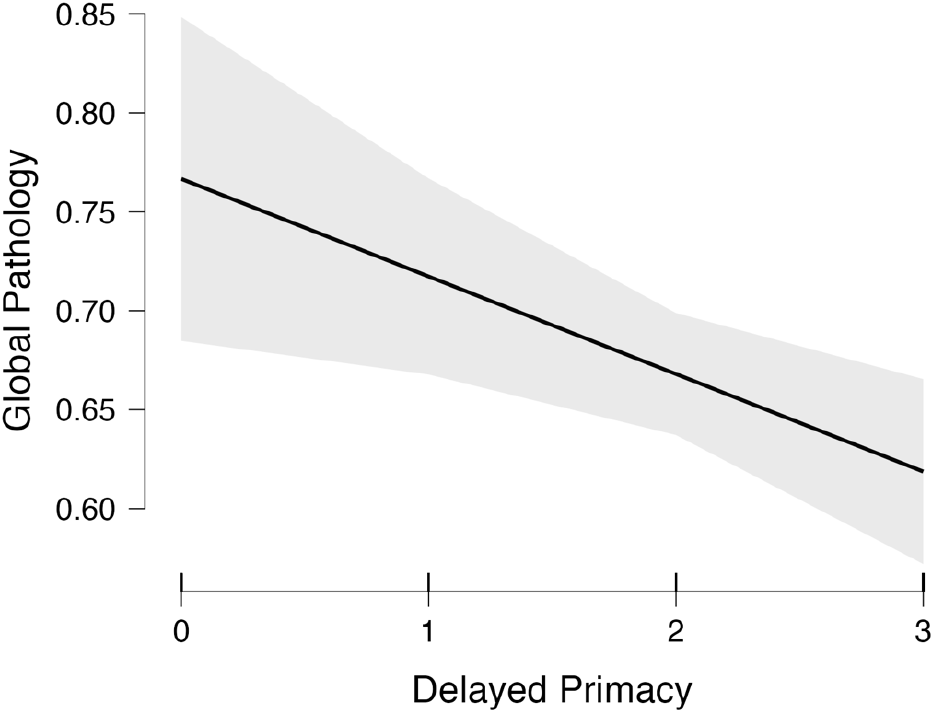
Marginal effects plot. Global Alzheimer’s disease pathology (y-axis) by delayed primacy (x-axis), and 95% confidence intervals.

Secondary analyses with neuritic plaques presented the same pattern of results, as this AD-related pathology was best predicted by delayed primacy, BF_10_ = 4.163, with moderate evidence. This model was over twice as strong as the second best model (BF_10_ = 1.560), again including only total delayed recall. The mean coefficient for delayed primacy was -0.041 (0.038; 95% credible intervals from -0.109 to <0.001), indicating that more neuritic plaque pathology was associated to poorer delayed primacy at baseline. BF_inclusion_ was 1.879, suggesting that adding delayed primacy to the model improved its fit to the data by 88%.

None of the models predicted diffuse plaques better than the null models including the covariates (best BF_10_ < 1). However, total delayed recall was an extremely strong predictor of neurofibrillary tangles (BF_10_ = 2891.180), narrowly beating out the model combining delayed primacy and immediate primacy (BF_10_ = 2592.755). The mean coefficient for total delayed recall was -0.022 (0.019; 95% credible intervals from -0.053 to 0), indicating that more neurofibrillary tangle pathology was associated to poorer delayed recall at baseline. BF_inclusion_ was 1.969, suggesting that adding delayed primacy to the model improved its fit to the data by 97%.

## 4. Discussion

In this secondary data analysis of over 1000 decedents who had participated in an RADC cohort study, we tested the hypothesis that a measure of delayed primacy, extracted from the CERAD memory test, would be effective in predicting *post mortem* AD-related pathology from a cognitively unimpaired baseline. These efforts are in line with the general goal of providing affordable, accessible and accurate tools to aid early detection and diagnosis of dementia of the Alzheimer’s type. We propose here a way to estimate delayed primacy in CERAD despite the fact that the word list order is changed over three consecutive learning trials: we suggest taking into account delayed recall performance for the words presented *first* in each of the three learning trials, i.e., the A1, B1 and C1 words. Our primary analysis showed that delayed primacy was the best predictor of global AD pathology, outperforming both total and delayed recall. In secondary analyses, delayed primacy was again the best predictor of neuritic plaques, while delayed recall was better for neurofibrillary tangles.

One observation from these results is that delayed recall measures, whether specifically for the primacy words or for the whole list, associated better with neuropathological outcomes than immediate recall measures. This finding is not surprising given the reported links between delayed recall and consolidation – as consolidation of new information requires time and biochemical changes in the central nervous system, delayed responses are more likely to be able to assess overall health of this mechanism than immediate responses [25,27,26,36].

However, primacy vs. whole list recall appeared to associate better with different types of post mortem pathology. Neuritic plaques, which are the consequence of amyloid-β protein aggregation, and neurofibrillary tangles, which derive from the accumulation of tau proteins, are the hallmarks of Alzheimer’s disease [37,38,39]. Delayed primacy was most sensitive to the former, while delayed recall was most sensitive to the latter. The association of delayed primacy with neuritic plaques is somewhat consistent with a previous report in a different dataset, showing that poor delayed primacy (and primacy forgetting) in story recall was associated with increased amyloid load, as measured with Pittsburgh compound-B PET imaging [40]. While the exact neurobiological mechanism linking delayed primacy recall to amyloid deposition is not clear, we have previously suggested that this association may be mediated by a failure to consolidate effectively the temporal information paired with the items, thus leading to poorer information clustering and, eventually, retention. This hypothesis, which requires further testing, is tentatively consistent with the observation that amyloid deposition in the brain begins in the neocortex [41], where associative memory function may be expressed [42]. An alternative, but not mutually exclusive, possibility is also that delayed primacy may be more sensitive to neuritic plaques because they will begin to accumulate years before neurofibrillary tangles, as per the amyloid cascade hypothesis [38], and therefore delayed primacy may be sensitive to early subtle changes that total delayed recall is unable to detect.

With regards to delayed recall and its preferential association with neurofibrillary tangles, we similarly do not have a specific mechanism to propose, except for highlighting how tangles have been observed to appear initially in the medial temporal region, first in the entorhinal cortex and then the hippocampus, which are areas strongly associated with memory consolidation [43]. To determine whether delayed recall performance broadly, as opposed to non-primacy delayed recall specifically, was preferentially associated with neurofibrillary tangles, we carried out a *post-hoc* analysis analogous to the main analyses. Outcome was *post mortem* neurofibrillary tangles; control variables were unchanged; and predictors were delayed recall, and the unstandardised residuals of regressing delayed primacy out of delayed recall (i.e., a measure of delayed recall independent of delayed primacy). This new analysis did not show primacy-independent delayed recall to be a good predictor of tangles, BF_10_ = 1.363, thus confirming the original conclusion that it was delayed recall *overall* that best associated with neurofibrillary tangles.

Another note is that clinical composite scores, such as the global cognitive function index we employed in our analyses (which, however, to note, was not norm-corrected), are more likely to be used in clinical settings than raw scores. Although composite scores have shown reduced variability and stronger associations with amyloid burden compared to raw scores in some instances [44,45], they have also been criticised for missing out on subtle cognitive changes [46]. In particular, in our analyses, the global cognitive function index, which included performance over 19 standardised neuropsychological tests, was not predictive of *post mortem* neuropathology in our analyses, and therefore underperformed when compared to the delayed primacy and delayed recall raw scores.

A clear limitation of this study is that the exact mechanisms underlying the link between delayed primacy, delayed recall, and other portions of the serial position, and AD-related pathology remain speculative at this stage. While a modest number of studies have examined the association between serial position performance and cognitive impairment, few have tackled the neurocognitive bases of this association. Further research, possibly involving functional neuroimaging and connectivity, should be considered towards this goal.

To conclude, both CERAD-derived delayed primacy and delayed recall predicted *post mortem* AD pathology in individuals who were classed as unimpaired at baseline, above and beyond demographics. These measures displayed different associations – delayed primacy correlated with global pathology and neuritic plaques, while delayed recall correlated with neurofibrillary tangles – and both performed better than the overall composite score. All in all, both the traditional recall scores and their process approach counterparts should be considered in research and clinical settings where the CERAD memory test is employed.

## References

[1] Olivari BS, French ME, McGuire LC. The Public Health Road Map to Respond to the Growing Dementia Crisis. Innovation in Aging 2020;4(1). https://doi.org/10.1093/geroni/igz043

[2] Bradford A, Kunik ME, Schulz P, Williams SP, Singh H. Missed and delayed diagnosis of dementia in primary care: prevalence and contributing factors. Alzheimer Dis Assoc Disord. 2009;23(4):306–314. https://doi.org/10.1097/wad.0b013e3181a6bebc

[3] Chodosh J, Petitti DB, Elliott M, Hays RD, Crooks VC, Reuben DB, Galen Buckwalter J, Wenger N. Physician recognition of cognitive impairment: evaluating the need for improvement. J Am Geriatr Soc. 2004;52(7):1051–1059. https://doi.org/10.1111/j.1532-5415.2004.52301.x

[4] Werner P, Karnieli-Miller O, Eidelman C. Current knowledge and future directions about the disclosure of dementia: a systematic review of the first decade of the 21st Century. Alzheimers Dement. 2013;9(2):e74.#x2013;88. doi:10.1016/j.jalz.2012.02.006

[5] Mattsson N, Carrillo MC, Dean RA, Devous Sr MD, Nikolcheva T, Pesini P, Salter H, Potter WZ, Sperling RS, Bateman RJ, Bain LJ. Revolutionizing Alzheimer’s disease and clinical trials through biomarkers. Alzheimer’s & Dementia: Diagnosis, Assessment & Disease Monitoring. 2015;1(4):412–9. https://doi.org/10.1016/j.dadm.2015.09.001

[6] Cullen NC, Leuzy A, Janelidze S, Palmqvist S, Svenningsson AL, Stomrud E, Dage JL, Mattsson-Carlgren N, Hansson O. Plasma biomarkers of Alzheimer’s disease improve prediction of cognitive decline in cognitively unimpaired elderly populations. Nature Communications. 2021;12(1):3555. https://doi.org/10.1038/s41467-021-23746-0

[7] Manera V, Rovini E, Wais P. Early detection of neurodegenerative disorders using behavioral markers and new technologies: New methods and perspectives. Frontiers in Aging Neuroscience. 2023;15. https://doi.org/10.3389/fnagi.2023.1149886

[8] Walker R, Paddick SM. Dementia prevention in low-income and middle-income countries: a cautious step forward. The Lancet Global Health. 2019;7(5):e538–9. https://doi.org/10.1016/s2214-109x(19)30169-x

[9] Albert MS, DeKosky ST, Dickson D, Dubois B, Feldman HH, Fox NC, Gamst A, Holtzman DM, Jagust WJ, Petersen RC, Snyder PJ. The diagnosis of mild cognitive impairment due to Alzheimer’s disease: recommendations from the National Institute on Aging-Alzheimer’s Association workgroups on diagnostic guidelines for Alzheimer’s disease. Alzheimer’s & dementia. 2011;7(3):270–9. https://doi.org/10.1016/j.jalz.2011.03.008

[10] De Simone MS, Perri R, Fadda L, Caltagirone C, Carlesimo GA. Predicting progression to Alzheimer’s disease in subjects with amnestic mild cognitive impairment using performance on recall and recognition tests. Journal of neurology. 2019;266:102–11. https://doi.org/10.1007/s00415-018-9108-0

[11] De Tollis M, De Simone MS, Perri R, Fadda L, Caltagirone C, Carlesimo GA. Verbal and spatial memory spans in mild cognitive impairment. Acta Neurologica Scandinavica. 2021;144(4):383–93. https://doi.org/10.1111/ane.13470

[12] Dubois B, Feldman HH, Jacova C, DeKosky ST, Barberger-Gateau P, Cummings J, Delacourte A, Galasko D, Gauthier S, Jicha G, Meguro K. Research criteria for the diagnosis of Alzheimer’s disease: revising the NINCDS–ADRDA criteria. The Lancet Neurology. 2007;6(8):734–46. https://doi.org/10.1016/s1474-4422(07)70178-3

[13] Mueller KD, D. L, Bruno D, Betthauser T, Christian B, Johnson S, Hermann B, Koscik RL. Item-level story recall predictors of amyloid-beta in late middle-aged adults at increased risk for Alzheimer’s disease. Frontiers in Psychology. 2022;3553. https://doi.org/10.3389/fpsyg.2022.908651

[14] Libon DJ, Swenson R, Ashendorf L, Bauer RM, Bowers D. Edith Kaplan and the Boston process approach. The Clinical Neuropsychologist. 2013;27(8):1223–33. https://doi.org/10.1080/13854046.2013.833295

[15] Milberg WP, Hebben N, Kaplan E, Grant I, Adams K. The Boston process approach to neuropsychological assessment. Neuropsychological assessment of neuropsychiatric and neuromedical disorders. 2009;3:42–65. https://doi.org/10.1093/arclin/act087

[16] Murdock Jr BB. The serial position effect of free recall. Journal of experimental psychology. 1962;64(5):482. https://doi.org/10.1037/h0045106

[17] Kelly MO, Risko EF. Offloading memory: Serial position effects. Psychonomic bulletin & review. 2019;26:1347–53. https://doi.org/10.3758/s13423-019-01615-8

[18] Neath I. Evidence for similar principles in episodic and semantic memory: The presidential serial position function. Memory & Cognition. 2010; 38(5), 659–666. https://doi.org/10.3758/mc.38.5.659

[19] Azizian A, Polich J. Evidence for attentional gradient in the serial position memory curve from event-related potentials. Journal of cognitive neuroscience. 2007;19(12):2071–81. https://doi.org/10.1162/jocn.2007.19.12.2071

[20] Rosen VM, Bergeson JL, Putnam K, Harwell A, Sunderland T. Working memory and apolipoprotein E: what’s the connection?. Neuropsychologia. 2002;40(13):2226–33. https://doi.org/10.1016/s0028-3932(02)00132-x

[21] La Rue A, Hermann B, Jones JE, Johnson S, Asthana S, Sager MA. Effect of parental family history of Alzheimer’s disease on serial position profiles. Alzheimer’s & Dementia. 2008;4(4):285–90. https://doi.org/10.1016/j.jalz.2008.03.009

[22] Kloth N, Lemke J, Wiendl H, Meuth SG, Duning T, Johnen A. Serial position effects rapidly distinguish Alzheimer’s from frontotemporal dementia. Journal of Neurology. 2020;267:975–83. https://doi.org/10.1007/s00415-019-09662-w

[23] Gicas KM, Honer WG, Wilson RS, Boyle PA, Leurgans SE, Schneider JA, Bennett DA. Association of serial position scores on memory tests and hippocampal-related neuropathologic outcomes. Neurology. 2020;95(24):e3303–12. https://doi.org/10.1212/wnl.0000000000010952

[24] Gicas KM, Honer WG, Leurgans SE, Wilson RS, Boyle PA, Schneider JA, Bennett DA. Longitudinal change in serial position scores in older adults with entorhinal and hippocampal neuropathologies. Journal of the International Neuropsychological Society. 2022;1–11. https://doi.org/10.1017/s1355617722000595

[25] Bruno D, Reiss PT, Petkova E, Sidtis JJ, Pomara N. Decreased recall of primacy words predicts cognitive decline. Archives of clinical neuropsychology. 2013;28(2):95–103. https://doi.org/10.1093/arclin/acs116

[26] Talamonti D, Koscik R, Johnson S, Bruno D. Predicting early mild cognitive impairment with free recall: The primacy of primacy. Archives of Clinical Neuropsychology. 2020;35(2):133–42. https://doi.org/10.1093/arclin/acz013

[27] Bruno D, Grothe MJ, Nierenberg J, Zetterberg H, Blennow K, Teipel SJ, Pomara N. A study on the specificity of the association between hippocampal volume and delayed primacy performance in cognitively intact elderly individuals. Neuropsychologia. 2015;69:1–8. https://doi.org/10.1016/j.neuropsychologia.2015.01.025

[28] Brueggen K, Kasper E, Dyrba M, Bruno D, Pomara N, Ewers M, Duering M, Bürger K, Teipel SJ. The primacy effect in amnestic mild cognitive impairment: associations with hippocampal functional connectivity. Frontiers in Aging Neuroscience. 2016;8:244. https://doi.org/10.3389/fnagi.2016.00244

[29] Bennett DA, Buchman AS, Boyle PA, Barnes LL, Wilson RS, Schneider JA. Religious orders study and rush memory and aging project. Journal of Alzheimer’s disease. 2018;64(1):S161–89. https://doi.org/10.3233/jad-179939

[30] Barnes LL, Shah RC, Aggarwal NT, Bennett DA, Schneider JA. The Minority Aging Research Study: ongoing efforts to obtain brain donation in African Americans without dementia. Current Alzheimer Research. 2012;9(6):734–45. https://doi.org/10.2174/156720512801322627

[31] Marquez DX, Glover CM, Lamar M, Leurgans SE, Shah RC, Barnes LL, Aggarwal NT, Buchman AS, Bennett DA. Representation of older Latinxs in cohort studies at the Rush Alzheimer’s Disease Center. Neuroepidemiology. 2020;54(5):404–18. https://doi.org/10.1159/000509626

[32] Bennett DA, Wilson RS, Schneider JA, Evans DA, Beckett LA, Aggarwal NT, Barnes LL, Fox JH, Bach J. Natural history of mild cognitive impairment in older persons. Neurology. 2002;59(2):198–205. https://doi.org/10.1212/wnl.59.2.198

[33] Bennett DA, Schneider JA, Aggarwal NT, Arvanitakis Z, Shah RC, Kelly JF, Fox JH, Cochran EJ, Arends D, Treinkman AD, Wilson RS. Decision rules guiding the clinical diagnosis of Alzheimer’s disease in two community-based cohort studies compared to standard practice in a clinic-based cohort study. Neuroepidemiology. 2006;27(3):169–76. https://doi.org/10.1159/000096129

[34] Wilson RS, Boyle PA, Yu L, Barnes LL, Sytsma J, Buchman AS, Bennett DA, Schneider JA. Temporal course and pathologic basis of unawareness of memory loss in dementia. Neurology. 2015;85(11):984–91. https://doi.org/10.1212/wnl.0000000000001935

[35] Morris JC, Mohs RC, Rogers H. Consortium to establish a registry for Alzheimer’s Disease (CERAD) clinical and neuropsychological. Psychopharmacology bulletin. 1989;24(3-4):641.

[36] Gomar JJ, Bobes-Bascaran MT, Conejero-Goldberg C, Davies P, Goldberg TE, Alzheimer’s Disease Neuroimaging Initiative. Utility of combinations of biomarkers, cognitive markers, and risk factors to predict conversion from mild cognitive impairment to Alzheimer disease in patients in the Alzheimer’s disease neuroimaging initiative. Archives of general psychiatry. 2011;68(9):961–9. https://doi.org/10.1001/archgenpsychiatry.2011.96

[37] Hampel H, Hardy J, Blennow K, Chen C, Perry G, Kim SH, Villemagne VL, Aisen P, Vendruscolo M, Iwatsubo T, Masters CL. The amyloid-β pathway in Alzheimer’s disease. Molecular psychiatry. 2021;26(10):5481–503. https://doi.org/10.1038/s41380-021-01249-0

[38] Hardy JA, Higgins GA. Alzheimer’s disease: the amyloid cascade hypothesis. Science. 1992;256(5054):184–5. https://doi.org/10.1126/science.1566067

[39] Tiraboschi P, Hansen LA, Thal LJ, Corey-Bloom J. The importance of neuritic plaques and tangles to the development and evolution of AD. Neurology. 2004;62(11):1984–9. https://doi.org/10.1212/01.wnl.0000129697.01779.0a

[40] Bruno D, Mueller KD, Betthauser T, Chin N, Engelman CD, Christian B, Koscik RL, Johnson SC. Serial position effects in the Logical Memory Test: Loss of primacy predicts amyloid positivity. Journal of neuropsychology. 2021;15(3):448–61. https://doi.org/10.1111/jnp.12235

[41] Thal DR, Rüb U, Orantes M, Braak H. Phases of Aβ-deposition in the human brain and its relevance for the development of AD. Neurology. 2002;58(12):1791–800. https://doi.org/10.1212/wnl.58.12.1791

[42] Aschauer D, Rumpel S. The sensory neocortex and associative memory. Behavioral Neuroscience of Learning and Memory. 2018;177–211. https://doi.org/10.1007/7854_2016_453

[43] Metaxas A, Kempf SJ. Neurofibrillary tangles in Alzheimer’s disease: elucidation of the molecular mechanism by immunohistochemistry and tau protein phospho-proteomics. Neural regeneration research. 2016;11(10):1579. https://doi.org/10.4103/1673-5374.193234

[44] Bransby L, Lim YY, Ames D, Fowler C, Roberston J, Harrington K, Snyder PJ, Villemagne VL, Salvado O, Masters CL, Maruff P. Sensitivity of a Preclinical Alzheimer’s Cognitive Composite (PACC) to amyloid β load in preclinical Alzheimer’s disease. Journal of clinical and experimental neuropsychology. 2019;41(6):591–600. https://doi.org/10.1080/13803395.2019.1593949

[45] Jonaitis EM, Koscik RL, Clark LR, Ma Y, Betthauser TJ, Berman SE, Allison SL, Mueller KD, Hermann BP, Van Hulle CA, Christian BT. Measuring longitudinal cognition: individual tests versus74–84. https://doi.org/10.1016/j.dadm.2018.11.006

[46] Bock JR, Russell J, Hara J, Fortier D. Optimizing Cognitive Assessment Outcome Measures for Alzheimer’s Disease by Matching Wordlist Memory Test Features to Scoring Methodology. Frontiers in Digital Health. 2021;161. https://doi.org/10.3389/fdgth.2021.750549

